# A low-dose immunotherapy targeting Fc-gamma Receptors and Heparan Sulfate Proteoglycan to impact myeloid cells and control tumor growth in cancers with varying immunosuppressive profiles

**DOI:** 10.1101/2025.08.07.665051

**Authors:** Nora Kakwata-Nkor Deluce, Elodie Creton, Alexandra Savatier, Honorine Lucchi, Laureen Bardouillet, Oscar Pereira Ramos, Philippe Berthon, Michel Léonetti

**Affiliations:** Blue Bees Therapeutics, Parc Club-Orsay Université, 21 rue Jean Rostand, 91400 Orsay, France; Département Médicaments et Technologies pour la Santé, Université Paris Saclay, CEA, INRAE, SPI, 91191 Gif-sur-Yvette, France

**Keywords:** Myeloid-targeted immunotherapy, Dendritic cell activation, Fc-gamma receptor, Heparan-Sulfate Proteoglycan, FcγR/HSPG-engaging immunotherapy, highly immunosuppressed tumors

## Abstract

**Background:** Myeloid cells play a central role in cancer-associated immunosuppression. Indeed, their accumulation and functional reprogramming attenuate effective immune responses and may facilitate tumor progression. To modulate the activity of the myeloid subsets, we chose to target Fcγ receptors (FcγRs), since all myeloid cell populations variably express FcγRs. As FcγRIIb provides an inhibitory signaling that might alter efficacy, we developed an immunotherapy, active at a low dose to limit binding via this low-affinity receptor while still interacting with higher-affinity FcγRs. We engineered a Fc-based fusion protein whose activity is potentiated by its ability to engage both FcγRs and a coreceptor, Heparan Sulfate Proteoglycan (HSPGs). We designed and fused a HSPG-ligand, named T54, to a human IgG1-Fc molecule to produce the Fc-T54 fusion protein.

**Methods:** Fc-T54 was produced using recombinant technologies. Binding characteristics were assessed using ELISA and flow-cytometric assays. Immune activity was investigated using cell-culture assays. Ability to affect tumor growth was investigated using four syngeneic tumor models with deserted to inflamed characteristics that differ in their sensitivity to immune checkpoint inhibitors (ICIs). Tumor microenvironment (TME) was analysed by flow cytometry.

**Results:** Compared to Fc, Fc-T54 demonstrates superior binding to low-affinity FcγRs and interacts more effectively with human leukocytes, including neutrophils, and B lymphocytes, as well as with monocytes and dendritic cells (DCs) within peripheral blood mononuclear cells. In activation assays Fc-T54 increases the number of monocytes/macrophages and B-lymphocytes, decreases neutrophil abundance, and enhances DC activation. Fc-T54 also demonstrates enhanced interaction with murine DCs accompanied by increased activation. Subcutaneous administration of low dose Fc-T54 - or its murine surrogate - significantly inhibits tumor growth in immune-deserted and immune-excluded mouse models, and synergizes with anti-PD-1 therapy in the immune-inflamed model. TME analysis in the MB49 bladder cancer model reveals that the immunotherapy decreases the proportion of granulocytic myeloid-derived suppressor cells while increasing CD8+ T-cells and natural killer cells, promoting a microenvironment more prone to tumor control.

**Conclusions:** This new FcγR/HSPG-engaging immunotherapy, administered via subcutaneous route, offers a novel approach to modulate the myeloid compartment and expand therapeutic options for ICI-resistant, deserted/excluded tumors, and for inflamed tumors when used in combination regimens.

## INTRODUCTION

Myeloid cells are key mediators of cancer immunosuppression, shaping a tumor-permissive microenvironment by inhibiting T-cells, secreting cytokines, and recruiting regulatory networks ^1,2^. Immunosuppressive myeloid populations include myeloid-derived suppressor cells (MDSCs) ^3^, tumor-associated macrophages (TAMs) polarized toward an immunosuppressive phenotype ^4^, and dysfunctional dendritic cells (DCs) that fail to prime antitumor immunity and expand regulatory T cells ^5^. Current therapies targeting these cells, such as CSF-1R inhibitors or CD40 agonists, can enhance the efficacy of checkpoint blockade by reprogramming TAMs or reducing MDSC infiltration. However, these approaches face limitations, including incomplete myeloid depletion, enrichment of resistant myeloid subsets, compensatory neutrophil populations, and adverse events ^6–9^. Therefore, novel therapeutic strategies that overcome these resistance mechanisms and elicit durable responses are needed, particularly in highly immunosuppressed cancers characterized by abundant MDSCs and resistance to current immunotherapies ^7,10,11^. This context prompted us to develop a new immunotherapy targeting myeloid cells.

All the myeloid cell populations that arise from progenitor cells express, in a variable manner, Fc gamma receptors (FcγRs) that can modulate their activity through mechanisms initiated upon binding of the Fc domain of IgGs ^12^. However, although FcγRs are targeted by numerous mono-, bi- and tri-specific antibodies developed to date ^13^, they have never been considered as a sole immunotherapeutic target. Essentially because this family ^14^ has activating receptors, such as FcγRI, FcγRIIa, FcγRIIIa, but also inhibitory receptors, mostly FcγRIIb, that could thwart the immunotherapeutic effect ^15,16^. However, and interestingly, FcγRIIb is the member of the family with the lowest affinity for antibodies, with a KD of approximately 1–3 × 10⁻⁵ M. As a result, a non-specific antibody or Fc fragment used at concentrations below the KD for FcγRIIb would predominantly interact with activating FcγRs on myeloid cells. Nevertheless, such proteins, when in their monomeric forms, are generally inefficient at triggering cell-signaling and, consequently, at modulating cellular activity ^17,18^. Therefore, they must be modified to make them appropriate myeloid-cell modulators.

Many soluble endogenous proteins, including cytokines, chemokines, and growth factors, modulate cell activity and numerous biological processes through engagement of a receptor and a coreceptor ^19^. This mechanism of dual interaction is also shared by proteins of infectious microorganisms that use it to disseminate and/or alter immune defense mechanisms ^20^. Heparan sulfate proteoglycans (HSPGs) are glycoproteins ubiquitously distributed on the cell surface that often play the role of coreceptor^21^.HSPG-binding proteins interact with the heparan sulfate (HS) groups of HSPGs via regions rich in basic residues ^22^. Previously, we showed that coupling of a basic-rich HSPG-ligand to a protein antigen (Ag) enables an approximately 100- to 1000-fold decrease in immune complex concentrations required to be presented by antigen-presenting cells (APCs) via FcγRs to specific T-cells ^23^. In addition, we found that the fusion of such a HSPG-ligand to proteic vectors targeting various receptors allows an increase of the immune response *in vitro* ^24,25^ and *in vivo* ^26^. These findings, which highlight the value of proteins capable of binding both a receptor and an HSPG coreceptor in specific immune responses, led us to investigate whether a fusion protein combining an IgG1-Fc fragment and an HSPG ligand, termed Fc-T54, could: (i) efficiently target FcγR-expressing immune cells, (ii) modulate their activity, and (iii) impact tumor growth. Notably, ICI therapies currently used in patients are predominantly effective in tumors with inflamed characteristics, and much less in those displaying deserted or excluded immune phenotypes ^27,28^. Therefore, we set out to evaluate whether Fc-T54 can affect tumor growth using a panel of four mouse syngeneic tumor models which vary in their sensitivity to ICIs and encompass a spectrum from deserted to inflamed phenotypes.

In this report, we first describe the production of Fc-T54. We then characterize its binding properties to FcγRs and heparan sulfates, as well as its interactions with both human and murine immune cells. Furthermore, we demonstrate the capacity of Fc-T54 to modulate immune cell activity and show that, when administered at low doses via the subcutaneous route, this fusion protein or its murine surrogate can inhibit tumor growth in syngeneic models with excluded or deserted immune phenotypes, and, when used in combination regimens, in inflamed tumors. We additionally show that Fc-T54 alters the TME in a manner that favors tumor control. Collectively, these findings indicate that Fc-T54 represents a novel strategy to modulate the myeloid compartment and broaden therapeutic options for ICI-resistant deserted/excluded tumors.

## MATERIALS AND METHODS

### Protein production

Fc-T54 and Fc were produced following the protocol for expression in Expi-HEK cells (Gibco Transfection Kit ExpiFectamine^TM^293). Briefly, Expi-HEK cells (3 × 10⁶ cells/ml) were transfected with a pcDNA™3.4 plasmid encoding either Fc-T54 or Fc and ExpiFectamine^TM^293 Reagent. After 16 hours incubation at 37°C, medium was added. After 4 additional days of incubation, supernatants were collected, filtered, and supplemented with a protease-inhibitor cocktail. After dilution in 0.05% Tween-PBS, proteins were immunopurified using a Protein A column (ProSep®-vA High-Capacity, Millipore). The eluate was neutralized with Tris-HCl (1 M, pH8), and were submitted to a gel filtration (AKTA HiLoad® 16/600 Superdex® 200pg, Cytiva). Proteins were analyzed by 4-12% SDS-PAGE gel. Proteins were stored at -20°C. An identical protocol was used to generate a mouse IgG2a-Fc fragment, named mFc, and a mouse Fc-T54 surrogate, designated mFc-T54, in which the Fc encodes a murine IgG2a.

### Binding to heparin

To assess Fc-T54 and Fc binding to heparin, microtiter plates were coated with heparin-albumin (Hep-Alb) (1 µg/ml in 0.1 M phosphate buffer pH7.2) and saturated with 0.3% BSA (200 µl/well). Then, plates were washed and serial dilutions of Fc-T54 and Fc (0.1 M phosphate buffer, pH7.4, containing 0.1% BSA) were added to the wells. After 1-hour RT, plates were washed and 100 µl of Goat Anti-Human Horseradish Peroxidase conjugate (GAH-PO) was added. After 30 minutes incubation, plates were washed, and 200 µl of 2,2′-azino-bis 3-ethylbenzothiazoline-6-sulfonic acid (ABTS) was added. Absorbance was measured at 415 nm.

### Binding to FcγRI, FcγRIIIa, FcγRIIa and FcγRIIb/c receptors

Microtiter plates were coated with avidin (1 µg/ml in 0.1 M phosphate buffer pH7.2) and saturated with 0.3% BSA (200 µl/well) or only coated with 0.3% BSA (300 µl/well).

To assess binding to FcγRI both sets of plates were washed and incubated with serial dilutions of biotinylated FcγRI for 1 hour RT. After washings, a fixed amount of Fc-T54 or Fc (100 nM) was added. After one hour RT the plates were washed, and GAH-PO was added. Thirty minutes later, the plates were washed and ABTS was added. Absorbance was measured at 415 nm. To eliminate non-specific binding, the optical signal from BSA-coated plates was removed from that from avidin-coated plates.

To assess binding to FcγRIIIa, FcγRIIa and FcγRIIb/c, serial dilutions of Fc-T54 or Fc proteins were pre-incubated with biotinylated-FcγRIIIa or FcγRIIa (30 nM) or FcγRIIb/c (100 nM) overnight at 4°C. Mixtures were then transferred to plates coated with avidin. After one hour, the plates were washed, GAH-PO was added and binding was assessed as for FcγRI.

To assess binding to heparin and FcγRI, serial dilutions of Fc-T54 or Fc were incubated on heparin-albumin- and BSA-coated wells. After 1-hour RT, plates were washed and biotinylated-FcγRI (20 nM) was added. One hour later, plates were washed, streptavidin-PO was added. Thirty minutes later, plates were washed, and 200µl ABTS substrate was added. Absorbance was measured at 415 nm.

To assess binding to heparin and FcγRIIIa, FcγRIIa and FcγRIIb/c, a fixed amount of biotinylated FcγRIIIa, FcγRIIa or FcγRIIb/c (100nM) was incubated with two different dilutions of Fc-T54 or Fc (100 and 300 nM for binding to FcγRIIIa; 300 and 1000 nM for binding to FcγRIIa and FcγRIIb/c) overnight at 4°C. Then, mixtures were transferred into heparin-albumin- and BSA-coated wells. After 1-hour RT, plates were washed and treated with GAH-PO and ABTS as for heparin/FcγRI assay.

### Binding to Human Leukocytes

To assess Fc-T54 and Fc binding on human leukocytes, whole blood was collected from healthy donors and subjected to red blood cell lysis. Then, leukocytes were seeded in plates (5×10⁵ cells/well). Next, biotinylated Fc-T54 or Fc (10 nM or 30 nM) were added. After 30 minutes at 4°C, cells were washed and stained with streptavidin-PE cells and a panel of lineage-specific antibodies to identify dendritic cells, monocytes, neutrophils, natural killer cells, B lymphocytes, and T lymphocytes. Cells were incubated for 30 minutes at 4°C before flow cytometry (Attune Flow Cytometer, Thermo Fisher Scientific) and analysis using FlowJo 10.9 software. Binding was quantified as the percentage of PE-positive cells within each leukocyte subpopulation.

### Binding to DCs

To assess Fc-T54 and Fc binding on DCs, JAWSII DCs were used (CRL-11904). Cells were seeded in plates (10^5^ cells/well) with or without serial dilutions of Fc-T54, Fc, T-54, or Fc + T54. After 30 minutes at 4°C, plates were washed and a fluorescent Goat anti-human IgG-Ab (GOAH-PE) was added. Thirty minutes later, plates were washed and cells analyzed by flow cytometry. Binding was quantified as the percentage of PE-positive cells within total live cells. To assess binding specificity, a fixed amount (100 nM) of Fc-T54 was incubated with or without a rat anti-mouse CD16/CD32 Ab (clone 2.4G2, 4 µg/mL), soluble HS (3 µM), or a mixture of CD16/CD32 Ab and HS, respectively.

### Activation of DCs and involvement of the syk-tyrosine kinase pathway

To assess the ability to activate DCs, cells were seeded in plates (10^5^ cells/well) with or without a fixed dilution (300 nM) of Fc-T54, Fc, T-54, or Fc + T54. After 24 hours at 37°C, cells were washed, labelled with an anti-CD69 Ab and analyzed by flow cytometry. To assess the involvement of the syk-tyrosine kinase pathway, JAWSII cells were seeded in plates (10^6^ cells/ml) and incubated for 30 minutes at 37°C with or without an inhibitor of the syk-tyrosine kinase pathway (R406, 300 nM). Then, Fc-T54 or Fc (300 nM) was added. After 24 hours at 37°C, cells were washed, labelled with an anti-CD69 Ab and analyzed by flow cytometry.

### Assessment of Fc-T54 activity *in vitro* using human leukocytes

To assess the ability of Fc-T54 to affect human immune cells, leukocytes from healthy donors’ whole blood were used. Leukocytes were seeded in plates (5.10^6^ cells/ml, 100 µL/wells, RPMI medium supplemented with 5% human albumin serum) with or without Fc-T54 or Fc (300 nM). After 3 days of incubation, cells were labelled with a panel of lineage-specific antibodies. Cell activation was assessed using the CD69 and CD86 activation markers. Cell acquisition and analysis were performed by flow cytometry.

### Antitumor efficacy

Female C57BL/6 mice aged of 7 weeks (Janvier Lab, Le Genest-Saint-Isle, France) were housed in the animal facility of CEA Saclay. Experiments were performed in accordance with French government and institutional guidelines. MB49 (Bladder Carcinoma; CVL_7076), B16F10 (melanoma; CRL_6475), MC38 (colon carcinoma cell line; CVCL_B288), PAN02 (pancreatic Adenocarcinoma; CRL_2553). MB49 (0.5 or 1 M cells), MC38 (1 M cells), Pan02 (3 M cells) and B16F10 (0.2 M cells) tumor cell lines were injected s.c. in mice. Three to seven days later, mice were randomized and injected s.c. with either mFc-T54, mFc, or Fc-T54 (0.02 nmol per mouse) or PBS (Control) once for MB49 and B16F10 models, or four times for Pan02 and MC38. The Anti-PD-L1 biosimilar Ab Avelumab (Ave) and the mouse anti-PD-1 Ab (clone RMP1-14) were injected intraperitoneally (1 nmol per mouse). Tumor volume was monitored using a caliper. Animals that reached a body score limit before the end of the experiment were euthanized and excluded from the tumor-size follow-up. Animals were euthanized at the end of the experiment or before if they reached ethical limits approved by the ethical committee.

### Analysis of tumor-infiltrating immune cells

Analysis of immune cells within the TME was evaluated from tumors of euthanized MB49 mice previously treated with PBS or Fc-T54. Tumors were harvested, mechanically disrupted and digested with collagenase IV (2 mg/ml). Tumors were then disrupted using a 10 ml syringe plunger and incubated at 37°C for 30 min. Collagenase activity was stopped by adding 5 ml RPMI medium + 10% FBS. Cells were filtered with 40 μm cell-strainer to obtain single-cell suspensions. Cells were then stained with a panel of lineage-specific antibodies to identify myeloid derived suppressor cells (MDSCs), DCs, NK cells and lymphocytes. Cell acquisition and analysis were performed by flow cytometry.

## RESULTS

### Production of a fusion protein with FcγR- and HS-binding ability

To assess whether targeting of both FcgRs and a coreceptor can modulate activity of immune cells expressing these receptors, we produced two recombinant proteins. The first, called Fc-T54, contains a Fc (or Fc), a six residues linker, and a Tat domain (T54) derived from the HIV transcriptional transactivator that has a heparan sulfate (HS) binding site^29^. The Tat domain differs from that described by D. Knittel and coll. ^26^ by the lack of its three C-terminal basic residues. The second protein, called Fc, contains the Fc and the linker domain. Fc-T54 and Fc were expressed in HEK cells, immunopurified on a protein A gel column and were submitted to size-exclusion chromatography to remove high-molecular weight oligomers (Figure 1A). Gel electrophoresis shows that Fc-T54 migrates in a predominant band higher than that corresponding to the Fc indicating that T54 is fused to Fc (Figure 1B). For Fc-T54 and Fc the molecular weights found (38 kDa and 32 kDa respectively) are higher than their theoretical masses (30321.5 Da and 26078.5 Da) indicating post-translational modifications during expression in HEK cells.

**Figure 1.**
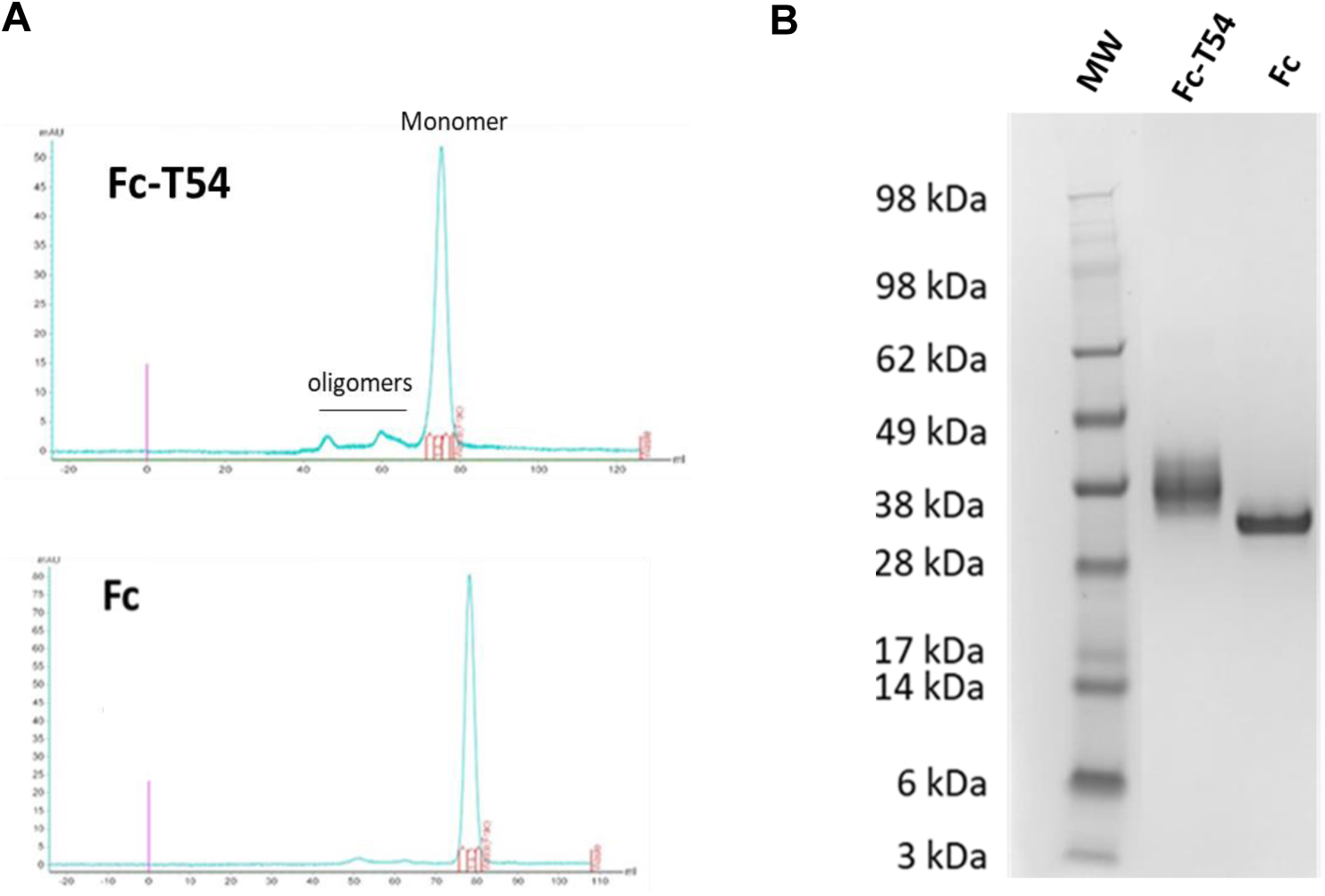
Characterization of purified Fc-T54 and Fc: integrity and homogeneity. **A:** Size-exclusion chromatography (SEC) profile of Fc-T54 and of Fc. Immuno-purified Fc-T54 and Fc were purified by SEC to remove oligomers. **B:** SDS-PAGE gel under denaturing conditions, showing markers of molecular weight (MW) and Fc-T54 and Fc after electrophoretic migration.

To determine whether the T54 domain endows the Fc with the ability to bind HS, Fc-T54 and Fc were compared by ELISA. Heparin, a sulfated polysaccharide representative of the HS-family, was used. No optical signal was detected for Fc, indicating that it does not bind heparin (Figure 2A). In contrast, Fc-T54 produced a dose-dependent optical signal, demonstrating its capacity to bind heparin.

**Figure 2.**
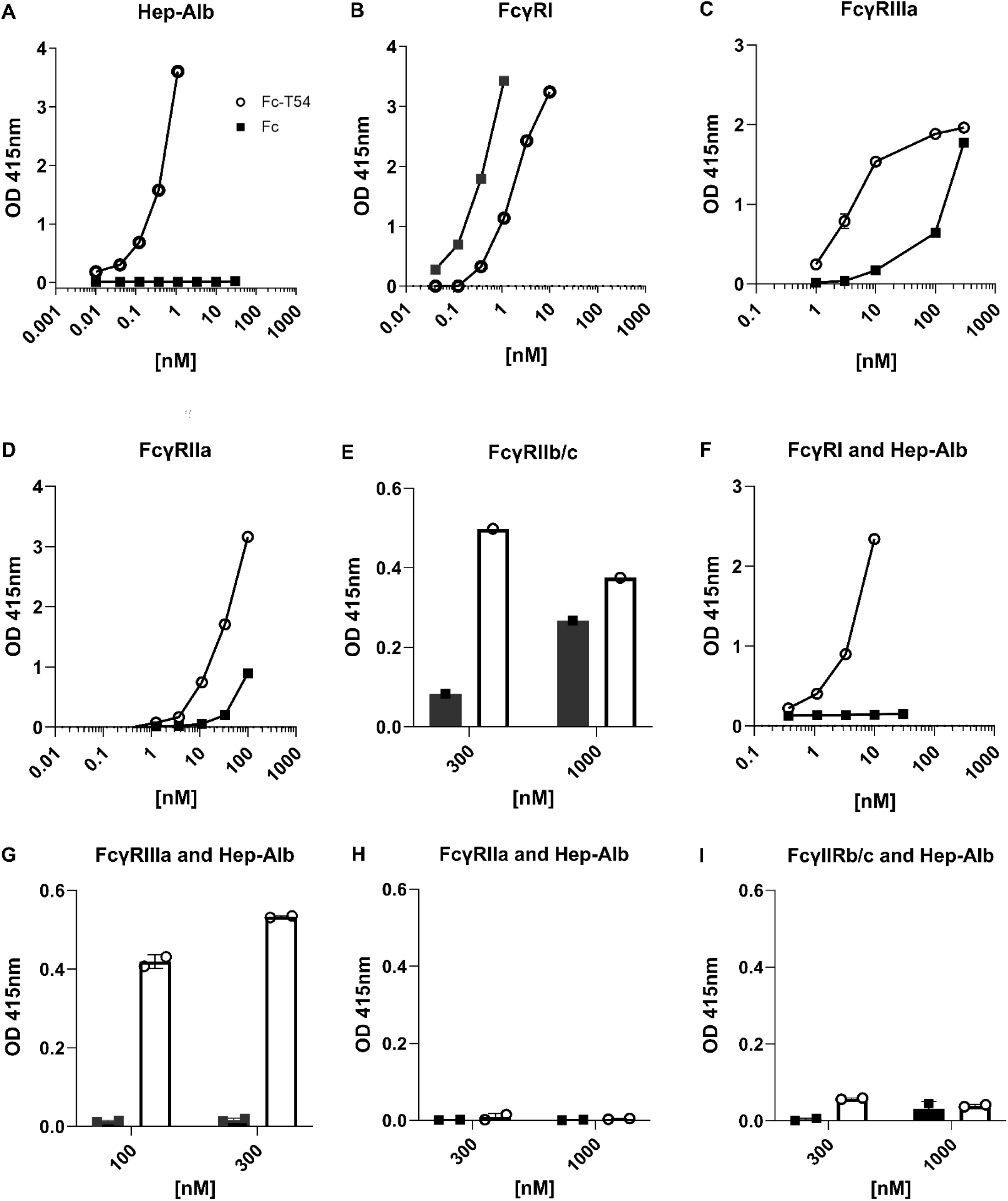
Binding of Fc-T54 and Fc to heparin and Fcg receptors. A: Binding of Fc-T54 and Fc proteins to heparin: Serial dilutions of Fc-T54 and Fc were incubated on heparin-albumin-coated plates. **B:** Binding to FcgRI: Serial dilutions of Fc-T54 and Fc were incubated on avidin-coated plates. Then, biotinylated-FcgRI was added. **C to E:** Binding to FcgRIIIa, FcgRIIa and FcgRIIb/c: Fc-T54 and Fc were pre-incubated overnight at 4°C with biotinylated-FcgRIIIa, -FcgRIIa and -FcgRIIb/c, respectively. Then, the mixtures were added to avidin-coated plates. Binding to the plates in **A-E** was detected using peroxidase-coupled goat anti-human IgG (PO-GOAHIgG) and ABTS as substrate. **F to I:** Study of simultaneous binding to Heparin and FcgRI, FcgRIIIa, FcgRIIa, FcgRIIb/c: **F:** Serial dilutions of Fc-T54 and Fc were incubated on heparin-albumin-coated plates, followed by an incubation with biotinylated FcgRI. **G to I:** Fc-T54 and Fc were pre-incubated with biotinylated-FcgRIIIa, -FcgRIIa, -FcgRIIb/c, respectively. Then, the mixtures were transferred to heparin-albumin-coated plates. Binding in **F to I** was detected using peroxidase-coupled streptavidin and ABTS as substrate.

### The HS-binding ability makes the Fc domain capable of interacting more efficiently with low-affinity human FcγRs and increases its ability to bind FcγR-expressing immune cells

To evaluate whether T54 conjugation affects the Fc ability to bind human Fcg receptors, Fc-T54 and Fc were assessed for their binding to soluble FcgRI, FcgRIIa, FcgRIIb, and FcgRIIIa. When mixtures of biotinylated FcgRI and serial dilutions of Fc-T54 or Fc were incubated on avidin-coated plates, a dose-dependent optical signal was observed (Fig. 2B). Notably, a lower concentration of Fc was required to achieve the same signal compared to Fc-T54, suggesting that covalent coupling of T54 to Fc reduces binding affinity for FcgRI. Both Fc-T54 and Fc also exhibited dose-dependent binding to FcgRIIIa. In this case, however, Fc-T54 required a lower concentration than Fc to reach the same signal, suggesting that T54 conjugation enhances binding to FcgRIIIa (Fig. 2C). A similar trend was observed for FcgRIIa and FcgRIIb, with increased binding seen for Fc-T54 compared to Fc. However, binding to these receptors was only detectable at higher concentrations for both Fc-T54 and Fc, consistent with the lower intrinsic affinity of the Fc domain for FcgRIIa and FcgRIIb ^12^.

Finally, we investigated the ability of Fc-T54 and Fc to bind both heparin and FcgRI, FcgRIIa, FcgRIIb, or FcgRIIIa. No optical signal was detected for Fc (Figure 2F to 2I), in agreement with its inability to bind heparin (figure 2A). A signal was measured for Fc-T54 with FcgRI or FcgRIIIa demonstrating its ability to bind these two receptors and heparin (figure 2F and 2G). For FcgRIIa and FcgRIIb the optical signal was too low to determine whether a simultaneous binding might occur.

Next, to assess whether T54 coupling also affects the ability of the Fc to bind to human FcgR-expressing immune cells, Fc-T54 and Fc were compared for their binding to human leukocytes from whole blood (figure 3). Fc and Fc-T54 did not interact with T-lymphocytes that do not express FcγRs (figure 3E). Fc also did not bind B-lymphocytes. In contrast, about 41% of the B-cells were bound by Fc-T54 suggesting that T54 coupling increases binding to FcγRIIb on these cells. The Fc was capable of binding NK cells, neutrophils, monocytes and DCs in agreement with the expression of FcγRs of highest affinity than FcγRIIb on the different cell types (figure 3A to 3D) ^12^. As compared to Fc, the proportion of neutrophils interacting with Fc-T54 was higher suggesting that T54 coupling increases binding to FcγRs on these cells. The difference between Fc and Fc-T54 was not significant for DCs and monocytes. For DCs, e assume that this might be related to their low amount (<1%) in leukocytes as compared to neutrophils (>50%) that would be mostly bound by the fusion protein. Interestingly, we found a significant difference between Fc and Fc-T54 for DCs and monocytes when using peripheral blood mononuclear cells (see figure S1). Surprisingly, while we observed a higher binding of the fusion protein to soluble FcγRIIIa by ELISA (see figure 2C), we did not find such a difference with NK cells that express FcγRIIIa. The reason for this behavior remains elusive.

**Figure 3.**
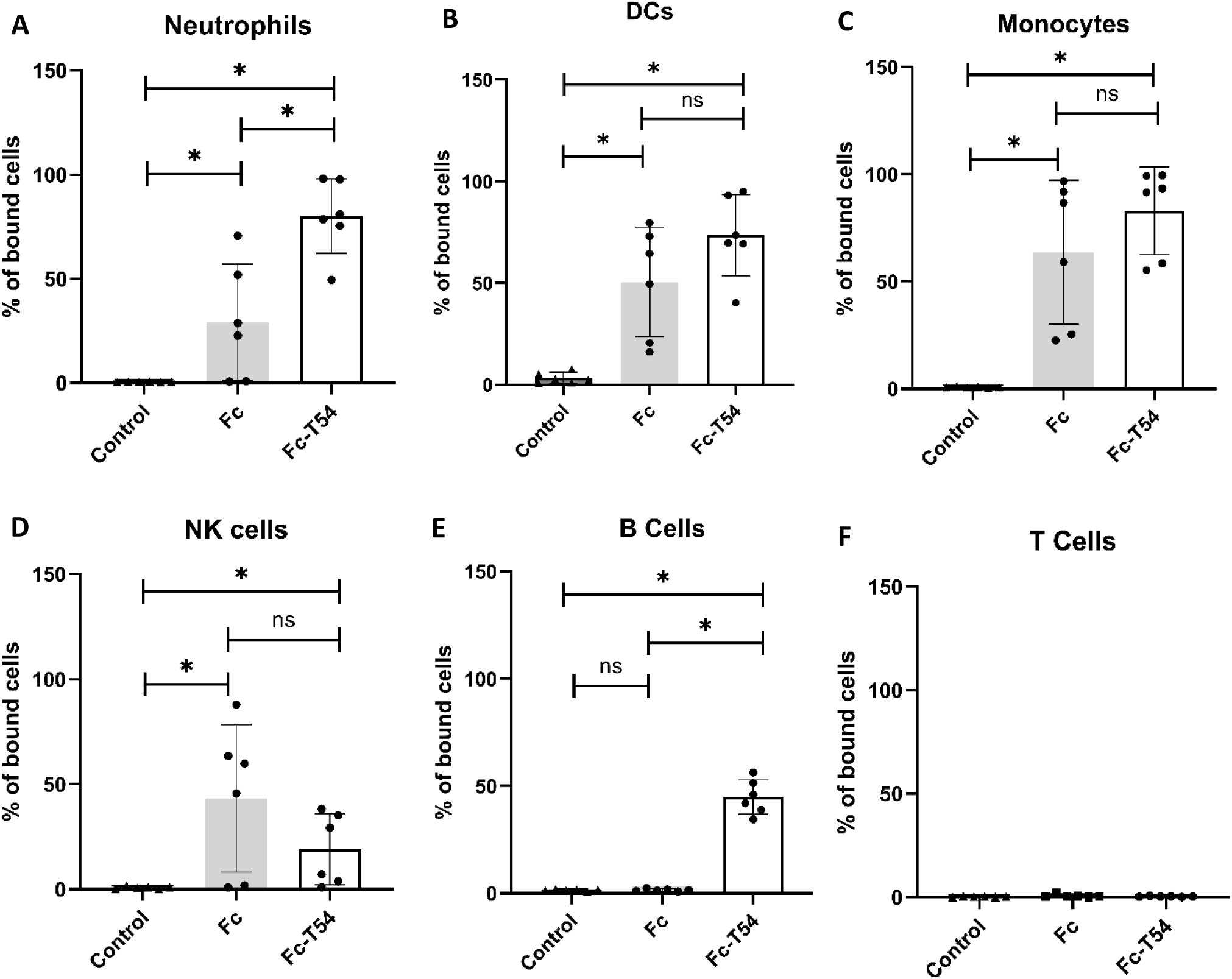
Binding of Fc-T54 and Fc to immune cells within leukocytes from whole blood. Leukocytes from human whole blood were incubated in the presence or absence of biotinylated Fc-T54 or Fc for 30 minutes at 4°C. After incubation, cells were washed and stained with a panel of lineage-specific antibodies to identify neutrophil, dendritic cells (DCs), monocytes, natural killer cells (NK cells), B lymphocytes (B cells), and T lymphocytes (T cells). Fc-T54 and Fc binding to these immune cell subsets was detected using fluorescent streptavidin. The percentage of fluorescent streptavidin-positive cells was plotted for each parent population. Data are presented as the mean ± SD of 3 independent experiences, paired t test, *p* < 0,05 (*), ns: non-significant.

### The Fc-T54 fusion protein modulates the activity of leukocytes from whole blood

To assess whether the fusion protein affects immune cells, we incubated *in vitro* leukocytes from human whole blood for three days in the presence or absence of either Fc-T54 or Fc. Then, we determined the proportion of the different subpopulations by flow cytometry. As shown in figure 4, Fc-T54 reduced the proportion of neutrophils, whereas Fc had no effect (figure 4A). As compared to Fc, the fusion protein increased the percentage of B lymphocytes and monocytes/macrophages while it had no effect on DCs, NK-cells and T-cells (figure 4B to 4F). To assess whether Fc-T54 could nevertheless impact the activity of the three latter populations we assessed the expression of i) the CD69 early activation marker on NK-cells, CD4+ and CD8+ T-cells, ii) the CD86 costimulatory molecule on DCs. We did not find an increased expression of CD69 on CD4+ T-cells (figure 4H). In contrast, as compared to the control, we found an upregulated CD69 expression on CD8+ T-cells (figure 4I) and NK-cells (figure 4J) as well as a higher CD86 expression on DCs (figure 4G) indicating that the fusion protein enhances activation of these immune cells.

**Figure 4.**
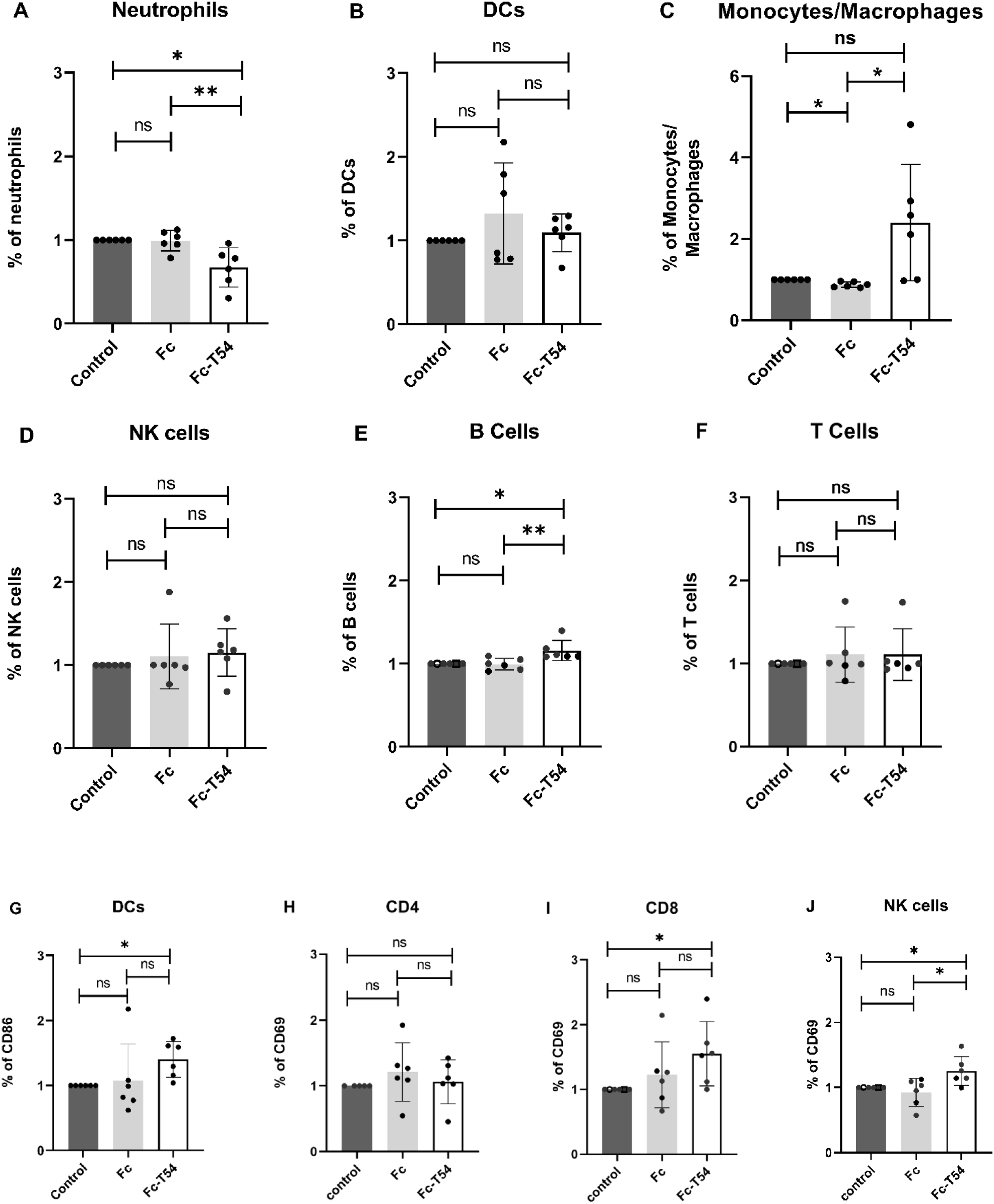
Fc-T54 impact on human immune cell subpopulations. Leukocytes from human whole blood were incubated for 3 days in the presence or absence of Fc-T54 or Fc. Cells were washed and stained with a panel of lineage-specific antibodies. The activation status was evaluated by quantifying CD69 expression for CD4+, CD8+ T-cells and NK-cells. CD86 expression was used for DCs. The percentage of cells and of activated cells was calculated relative to the total population of each cell type. Data are shown as mean ± SD of 3 independent experiences, paired t test, *p* < 0,05 (\**), p < 0,01* (**), ns: non-significant.

### The Fc-T54 fusion protein interacts more efficiently than Fc with mouse dendritic-cells via a mechanism involving FcγR-mediated binding and HS-binding ability

As human IgG1 can recognize mouse FcgRs ^30^, to validate the use of Fc-T54 in murine models we assessed its ability as well as that of Fc to bind to mouse FcgR-expressing immune cells. For this, we used a mouse DC-line, named JAWSII. We found that Fc or a mixture of Fc and T54 peptide binds less than 20% of DCs while Fc-T54 interacts with around 90% of cells (figure 5A). Next, to assess whether this enhanced binding depends on FcgR-binding and HS-binding ability of T54 we evaluated the impact of an anti-FcgRII/III ab (2.4G2 Ab) and of soluble HS on Fc-T54 binding to JAWSII cells. We found an about 30% lower interaction in the presence of the 2.4G2 Ab or HS (figure 5B) indicating that FcgR and HS-binding contribute to Fc-T54 binding. In addition, we observed about 80% inhibition when anti-FcgRII/III ab and HS were incubated together indicating a potentiated impact on binding. Altogether, these results demonstrate that Fc-T54 fusion protein interacts more efficiently than Fc with mouse DCs and that this increased interaction proceeds via a mechanism involving low-affinity FcgRs and HS-binding ability.

**Figure 5.**
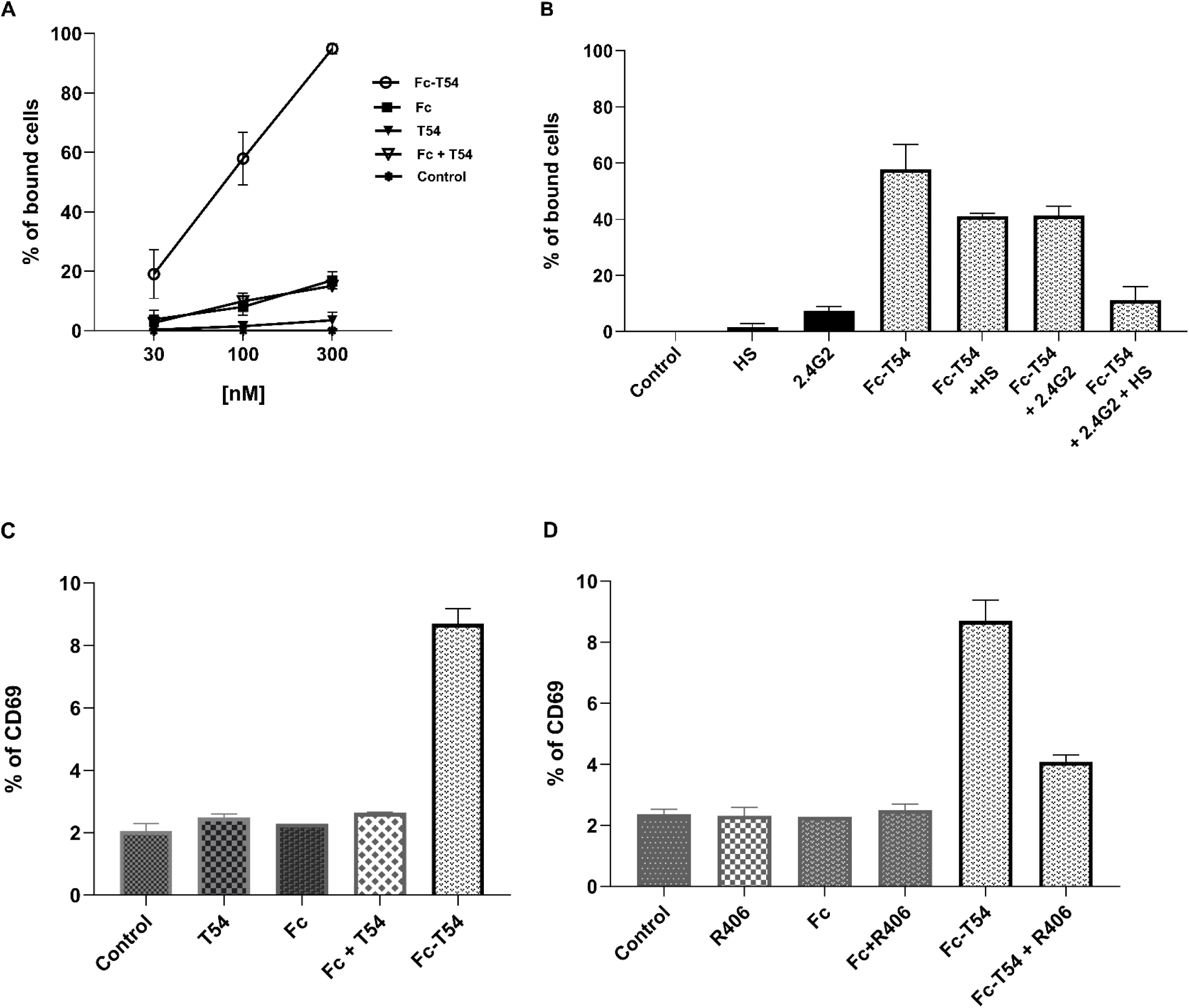
Fc-T54 ability to bind and activate mouse DCs. **A:** Fc-T54 and Fc binding to DCs. JAWSII DCs were incubated with Fc-T54, Fc, T54 or Fc + T54 for 30 min in 4°C. Then, cells were labeled with a fluorescent anti-human IgG Ab, and binding was assessed by flow cytometry. The percentage of fluorescent cells was calculated relative to the total number of live cells. **B:** Inhibition of Fc-T54 binding to DCs. Fc-T54 was incubated alone or in the presence of a Fc receptor-blocking Ab (2.4G2), soluble heparan sulfate (HS), or a combination of 2.4G2 and HS. Binding was analyzed as in A. **C.** To assess Fc-T54-mediated activation of DCs, JAWSII cells were incubated with or without Fc-T54, Fc, Fc + T54, and T54. After 24 hours, cells were labeled with a fluorescent anti-CD69 Ab. The percentage of CD69-positive cells was plotted relative to the number of live cells. **D:** DCs were pretreated with the syk pathway inhibitor R406. Thirty minutes later, cells were incubated with or without Fc-T54 and Fc. After 24 hours, cell activation was assessed as in C.

### The Fc-T54 fusion protein activates mouse dendritic cells more efficiently than Fc, through a mechanism that involves the Syk tyrosine kinase signaling pathway

To assess whether T54 covalent coupling to Fc increases the ability to trigger mouse DC activation, we assessed CD69 expression by the JAWSII murine DC-line after 24 hours incubation with Fc-T54, T54, Fc and Fc + T54, respectively. We found a tiny impact, if any, on this early activation marker with the molecules, i.e. T54, Fc and Fc + T54 (Figure 5C). In contrast, we observed a four-fold higher expression in the presence of Fc-T54. Therefore, these results indicated that T54 covalent coupling increases Fc ability to trigger mouse DC activation. Next, as FcgR-mediated activation proceeds through the Syk-tyrosine kinase pathway ^31^, we investigated its contribution using a Syk-pathway inhibitor, named R406. We incubated the fusion protein in the presence or absence of R406 and assessed CD69 expression. We found about a 70% decrease in Fc-T54-mediated CD69 expression in the presence of R406 indicating the contribution of the Syk-signaling pathway in activation (figure 5D). Altogether, these results demonstrate that T54 covalent coupling to Fc indeed increases the ability to trigger mouse DC activation and that this increased activation proceeds via a mechanism involving the Syk-signaling pathway.

### Subcutaneous administration of low doses of either a mouse Fc-T54 surrogate or Fc-T54 effectively controls the growth of tumors exhibiting diverse immunosuppressive profiles

In an initial series of experiments, to avoid biases associated with potential induction of anti-drug antibodies by Fc-T54 or differences in binding to mouse FcgRs^30^, we assessed tumor control efficacy using a surrogate for Fc-T54, termed mFc-T54, in which the human Fc domain was replaced with a murine IgG2a Fc. To validate the use of this surrogate, we performed preliminary experiments demonstrating that mFc-T54 interacts with and activates JAWSII dendritic cells more efficiently than mFc alone (Supplementary Figure S2). Then, we evaluated the effect of low doses subcutaneous injections of mFc-T54 (20pmol/injection) in four syngeneic tumor models.

The first model used was MB49 bladder cancer, which displays ‘desert’ characteristics, featuring a low proportion of T lymphocytes and a high proportion of myeloid-derived suppressor cells (MDSCs) ^32^, yet can still be controlled by standard anti-immune checkpoint antibodies ^33^. In this model, mFc-T54 inhibited tumor growth, while mFc and T54 alone were ineffective, indicating that only the fusion protein produces the antitumor effect (Figure 6A). Next, we employed a second ‘desert’ model: the highly aggressive B16F10 melanoma, characterized by limited immune cell infiltration ^34^. We observed a significant impact on tumor growth at D2 and D4 after mFc-T54 injection (Figure 6B). Here, mFc-T54 treatment significantly inhibited tumor growth at days 2 and 4 post-injection (Figure 6B). However, this effect did not persist over time (data not shown), suggesting that the treatment is primarily effective during the early tumorigenic phase in this highly immunosuppressed model. We also tested the Pan02 pancreatic tumor model, which displays ‘excluded’ features with immune infiltration at the tumor periphery, a high abundance of immunosuppressive cells (Tregs, MDSCs, and TAMs), and low CD8+ T cell content ^35^. In this model, mFc-T54 treatment significantly reduced tumor growth (Figure 6C). Finally, we evaluated the MC38 colorectal cancer model, which exhibits ‘inflamed’ characteristics with high immune cell infiltration and a low proportion of immunosuppressive cells ^36,37^. In this model, mFc-T54 treatment had no discernible effect on tumor growth (Figure 6D).

**Figure 6:**
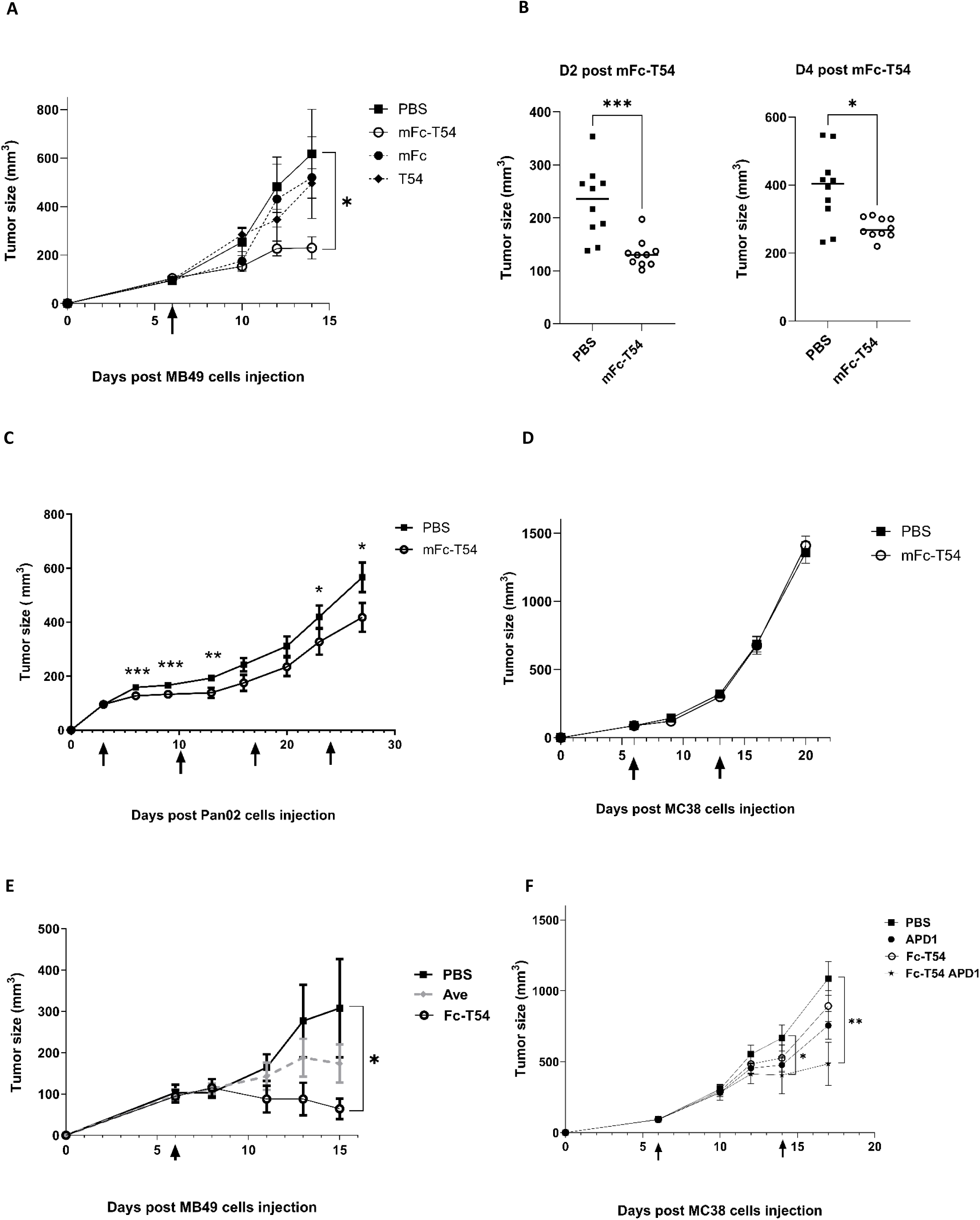
mFc-T54 and Fc-T54 reduce tumor growth in different syngeneic tumor models. **A:** Four groups of C57BL/6 mice (n=7 to 9 per group) were injected subcutaneously (s.c.) with 1× 10^6^ MB49 cells. Six days later, the groups were injected s.c. with mFc-T57, mFc, T54 and PBS, respectively. **B:** Two groups of C57BL/6 mice (n=10 per group) were injected s.c. s.c. with 0.2 × 10^6^ B16F10 cells. Ten days later, one group received mFc-T54, another received PBS. Tumor volume at days two and four is presented. **C:** Two groups of C57BL/6 mice (n=10 per group) were injected s.c. with 3 × 10^6^ Pan02 cells. Three, ten, seventeen and twenty-four days later, one group received mFc-T54, another received PBS. Tumor volume at days two and four is presented. **D:** Two groups of C57BL/6 mice (n=10 per group) were injected subcutaneously with 1 × 10^6^ MC38 cells. Six and thirteen days later, one group received mFc-T54, another received PBS. **E:** Three groups of C57BL/6 mice (n=5 per group) were injected s.c. with 0.5 × 10^6^ MB49 cells. Six days later, one group received Fc-T54, another received the Avelumab anti-PD-L1 biosimilar Ab, and the third received PBS. **F:** Four groups of C57BL/6 mice (n=5 to 8 per group) were injected subcutaneously with 0.5 × 10^6^ MC38 cells. Six and fourteen days later, one group received Fc-T54 s.c., another received the anti-PD-1 Ab i.p., the third Fc-T54 and anti-PD-1 Ab, the fourth received PBS. **A to F:** The arrow(s) indicate the day(s) of protein injection. Tumor growths were monitored every two days using a caliper. Mice were euthanized at the end of the experiment or when they reached the protocol-ethical defined limits. Data are presented as the mean ± SEM. Mann-Whitney test, *p* < 0,05 (\**), p < 0,01 (*\*\**), p < 0,001 (*\*\*\*).

Next, we used the MB49 model to evaluate the effect of a low-dose (20 pmol) subcutaneous injection of human Fc-T54, in comparison to an anti-immune checkpoint antibody administered at a standard dose for mice (100 µg, 667 pmol). To enable this comparison, we leveraged the ability of human anti-PD-L1 antibody drugs to cross-react with mouse PD-L1 ^38^. We selected Avelumab, commonly used in bladder cancer patients, and employed an Avelumab (Ave) biosimilar for the study. As shown in Figure 6E, Fc-T54 treatment led to a reduction in tumor size, and this effect was greater than that achieved with a 33-fold higher dose of Ave.

Finally, we assessed the effect of Fc-T54 in the MC38 model. Given the sensitivity of MC38 to immune checkpoint blockade ^39^ we also tested whether our immunotherapy could synergize with anti-PD-1 antibody treatment. In this inflamed tumor context, neither Fc-T54 nor anti-PD-1 antibody monotherapy significantly impacted tumor growth (Figure 6F). However, the combination treatment resulted in a marked reduction in tumor size, indicating a synergistic therapeutic benefit.

### Fc-T54 injection changes the pattern of immune cells infiltrating tumor microenvironment

To assess whether the effect of Fc-T54 on tumor growth is related to a change in the profile of immune cells infiltrating the TME, we selected the MB49 syngeneic model that has a TME with a significant proportion of MDSCs, DCs, T- and NK-cells ^32^. Given Fc-T54 injection into mice implanted with the amount of MB49 cells commonly used by research groups (i.e. 0.5 M) caused a reduction in tumor size (see figure 6D) that did not allow to collect enough cells for an accurate immunophenotyping, we injected a two-fold higher amount of cells to get a lower tumor-control. These conditions fit well since mice injected with Fc-T54 control tumor growth without size decrease (figure 7.A). Therefore, we collected tumors from the control and treated groups for immunophenotyping. As shown in Figure 7.B, as compared to the control group, the TME of animals injected with Fc-T54 exhibited a four time less amount of MDSCs, an unsignificant increase of DC, as well as an around three times increase in T- and NK-cells. These data therefore indicate that the fusion protein makes the TME more prone to tumor control.

**Figure 7:**
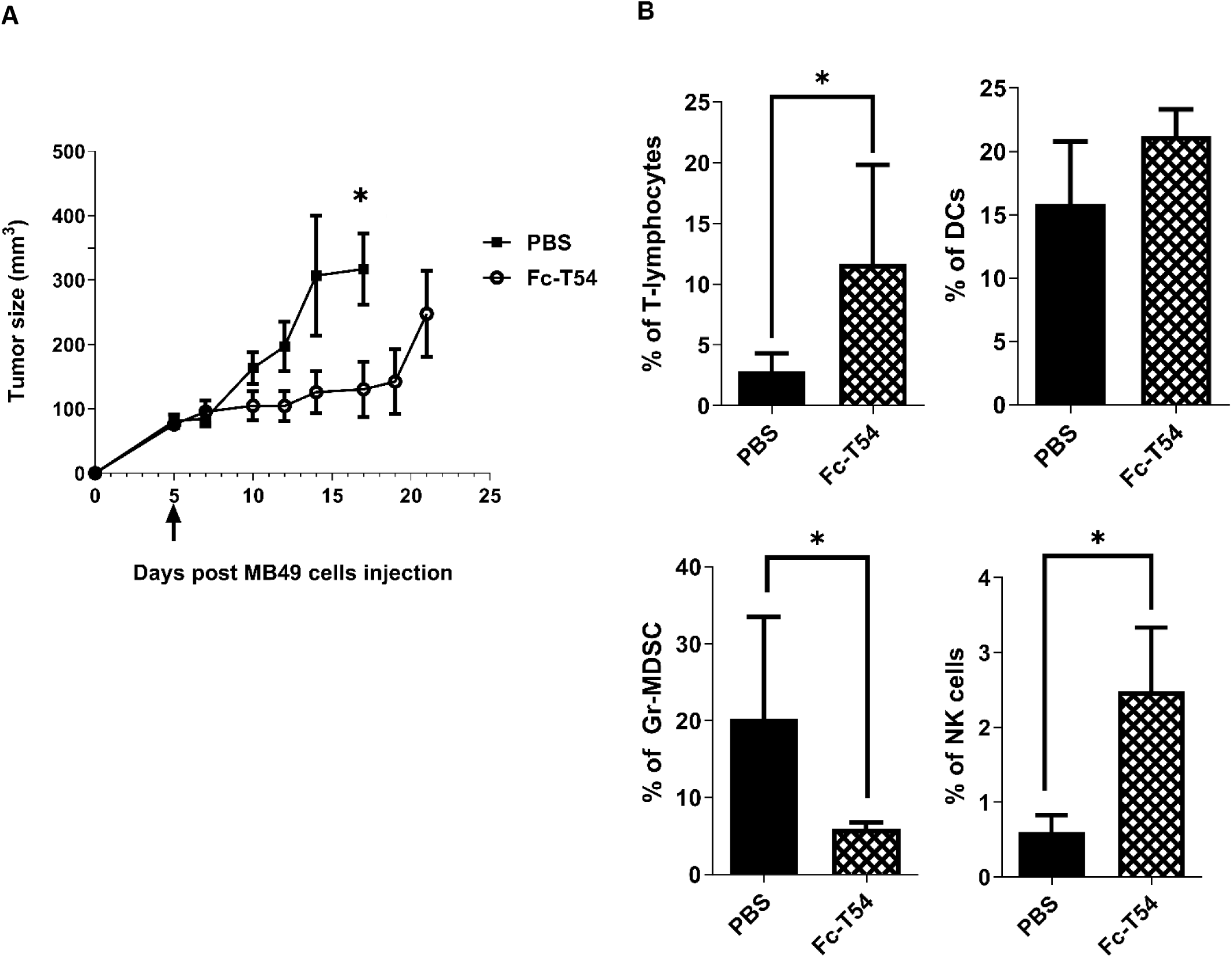
Fc-T54 injection changes the pattern of immune cells infiltrating MB49 tumor microenvironment. **A:** Mice injected with MB49 cells exhibit slower tumor growth when subsequently treated with Fc-T54. Two groups of C57BL/6 mice (n=5 and 6 per group) were injected s.c. with 1 M MB49 cells. Five days later, one group was injected with Fc-T54, another with PBS as control. Tumor growth was monitored using a caliper. **B:** Distinct immune cells infiltration in MB49 injected mice treated with Fc-T54 or PBS. Tumors were collected from mice injected with PBS or Fc-T54, and cells were labelled with fluorescent specific antibodies for T-lymphocytes, DCs, Gr-MDSC-cells and NK-cells, respectively. The proportion of each population is represented among total live cells. Data are presented as the mean ± SEM. Mann-Whitney test, *p < 0.05.

## DISCUSSION

We engineered our Fc-T54 fusion protein with the aim of modulating myeloid cell activity through the coengagement of FcgRs and heparan sulfate proteoglycans (HSPGs). As compared to Fc, Fc-T54 exhibits reduced binding to FcgRI and enhanced interaction with low-affinity FcgRIIa, FcgRIIb, and FcgRIIIa. Previous structural studies indicated that complex formation between the Fc moiety of an IgG1 and the different FcgRs are closely related with significant contributions from the lower hinge region and the CH2 domains. However, the nature and strength of these interactions vary due to distinct structural features, such as the presence of a shorter FG loop in the D2 domain of FcgRI and the influence of glycosylation on immunoglobulin CH2 domain, FcgRIIs and FcgRIIIa ^40^. Since Fc-T54 is a weaker binder than Fc to high affinity FcgRI and a stronger binder to low affinity FcgRs, we hypothesize that C-terminal fusion of T54 to Fc may induce long-range structural effects in the CH2 domains thereby differentially modulating affinity depending on the FcgR subtype. Coupling of T54 enables Fc-T54 to bind heparin, a representative molecule of the HS family. Thus, it makes the fusion protein capable of simultaneously interacting with both FcgRs and heparin. This dual-binding capability suggested that Fc-T54 could serve as a relevant molecule for simultaneously engaging FcgRs and HSPG coreceptors on immune cells expressing FcgRs, particularly myeloid cells.

Experiments performed with leukocytes from human whole blood show that Fc-T54 has a pattern of cell-binding similar to that of Fc, strongly suggesting that its interaction is primarily mediated by binding to FcγRs. Compared to Fc, the fusion protein interacts more strongly with the predominant myeloid population, i.e. neutrophils, indicating that coupling with the T54 domain enhances binding. Fc-T54 does not exhibit a significantly increased binding to monocytes, NK-cells, NKT-cells, or DCs. For the latter population, this may be due to its low proportion relative to neutrophils, as we found a significantly higher number of DCs bound by Fc-T54 as compared to Fc when using human PBMCs which are depleted of neutrophils (supplementary figure S1). Furthermore, we observed that the fusion protein binds more effectively than Fc to a murine DC-line, and this enhancement depends on FcgRII/III and HS-binding capacity. Collectively, these data indicate that Fc-T54 has an enhanced ability to bind FcgR-expressing myeloid cells.

In vitro, Fc-T54 affects the immune cells to which it binds more strongly. Incubation with Fc-T54 reduces the proportion of neutrophils, whereas Fc has no effect, suggesting the induction of inhibitory signals. The underlying mechanisms remain to be clarified, as these cells express activating FcgRIIa, FcgRIIc, and FcgRIIIb, and to a lesser extent, the inhibitory FcgRIIb ^14^. Fc-T54 increases the proportion of B lymphocytes and macrophages increases, indicating that it may induce activating signals. Blood dendritic cells (DCs) show upregulated CD86 expression, reflecting enhanced activation. Increased activation is also observed in a murine DC line. This enhancement is abrogated by a Syk tyrosine kinase pathway inhibitor, confirming the involvement of the FcgR signaling cascade ^31^. Collectively, these findings support the hypothesis that activation is modulated through the co-engagement of FcgRs and an HSPG co-receptor, thereby amplifying FcgR-mediated cell signaling.

The enhanced modulatory activity resulting from FcgR/HSPG co-receptor engagement prompted us to test low-dose Fc-T54 (20 pmoles, 30-60 times than typical ICIs doses ^41^) in multiple murine syngeneic tumor models. Given its ability to activate DCs, we prioritized the subcutaneous route due to the high density of DCs in the skin ^42,43^. This approach proved effective as Fc-T54 induced tumor control in three models. These findings support low-volume subcutaneous injection of Fc-T54 in clinical trials, an approach generally avoided for ICIs due to high-dose requirements. However, recent studies show that subcutaneous ICI administration can match intravenous efficacy while enhancing patient comfort and quality of life ^44,45^.

Certain ICIs can reshape the TME by increasing T cell infiltration and reducing immunosuppressive cells such as Tregs and MDSCs, thereby converting it into a more immunologically active milieu ^46^. Targeting MDSCs or reprogramming TAMs toward a pro-inflammatory phenotype enhances antitumor immunity ^47,48^. The activation and recruitment of DCs are also key factors in initiating and sustaining antitumor immune responses ^49^. Collectively, these studies illustrate that modulating the immune cell composition of the TME is a validated therapeutic avenue for controlling tumor growth^50^. In line with these findings, our Fc-T54 immunotherapy is capable of remodeling the TME of the MB49 syngeneic model ^32^ by decreasing the proportion of MDSCs and increasing its T-lymphocyte and NK-cell content. Thus, Fc-T54 transforms an immunosuppressive niche into one more amenable to tumor control.

Fc-T54 displays broad and versatile antitumor efficacy across syngeneic mouse models representing immune-desert, immune-excluded, and immune-inflamed tumor phenotypes. In these desert models, Fc-T54 overcomes pronounced local immunosuppression, notably controlling tumor growth in MB49 —where anti-PD-L1 is effective— and delaying early progression in the highly aggressive, rapidly growing B16F10 model, which is resistant to checkpoint inhibition. In the immune-excluded Pan02 model, Fc-T54 reduces tumor burden, suggesting its capacity to bypass exclusion barriers. In contrast, Fc-T54 alone has no effect in the immune-inflamed MC38 model, but combines synergistically with anti-PD-1 therapy. Altogether, these findings highlight Fc-T54 as a flexible immunotherapeutic option for tumors with diverse immunosuppression, growth dynamics, and ICI responsiveness, potentially expanding treatment avenues for ICI-resistant, desert/excluded tumors, and in combination regimens for inflamed tumors.

## Acknowledgements

We thank Moutuaata Mohamed Moutuou for her help in *in vitro* experiments; François Becher for mass spectrometry analysis.

## Contributors

NKND designed and performed the experiments and wrote the manuscript. EC, AS and HL performed the experiments in vitro and in animals. LB performed in vitro experiments and FACS analysis. OPR designed protein constructs and approaches for protein engineering. PB designed experiments and supervised part of the project. ML supervised the entire project, designed the experiments and revised the manuscript. All authors have read and agreed to the published version of the manuscript. NKND, PB and ML act as guarantors for this study.

## Ethics approval

All animal studies were performed in compliance with the Ethics Committee of CEA (CETEA DSV – comité n° 44). Blood samples were collected by “Etablissement Français du Sang” in compliance with its Ethics Committee.

