## Supplemental data 1 and 2 for "A low-dose immunotherapy targeting Fc-gamma Receptors and Heparan Sulfate Proteoglycan to impact myeloid cells and control tumor growth in cancers with varying immunosuppressive profiles"

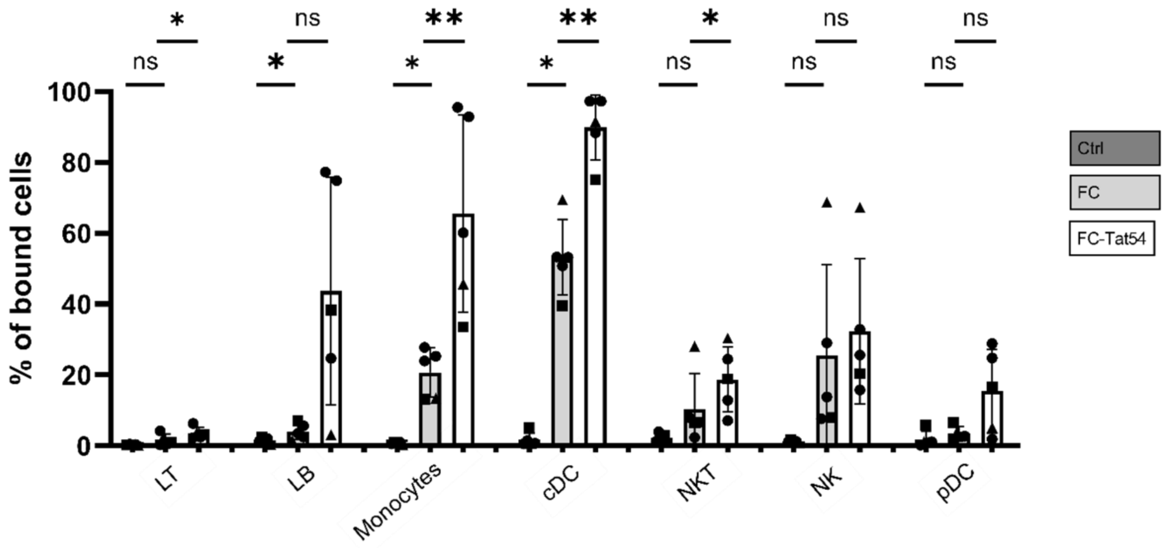


**Figure S1. Binding of Fc-T54 and Fc to immune cells within PBMCs.** PBMCs from human whole blood were incubated in the presence or absence of biotinylated Fc-T54 or Fc for 30 minutes at 4°C. After incubation, cells were washed and stained with a panel of lineage-specific antibodies to identify neutrophils, dendritic cells (DCs), monocytes, natural killer cells (Nk cells), B lymphocytes (B cells), and T lymphocytes (T cells). Fc-T54 and Fc binding to these immune cell subsets was detected using fluorescent streptavidin. The percentage of fluorescent streptavidin-positive cells was plotted for each parent population. Data are presented as the mean ± SD of 5 independent experiences, paired t test, *p* < 0,05 (*), *p < 0,01 (**), p < 0,001 (*****), ns: non-significant.

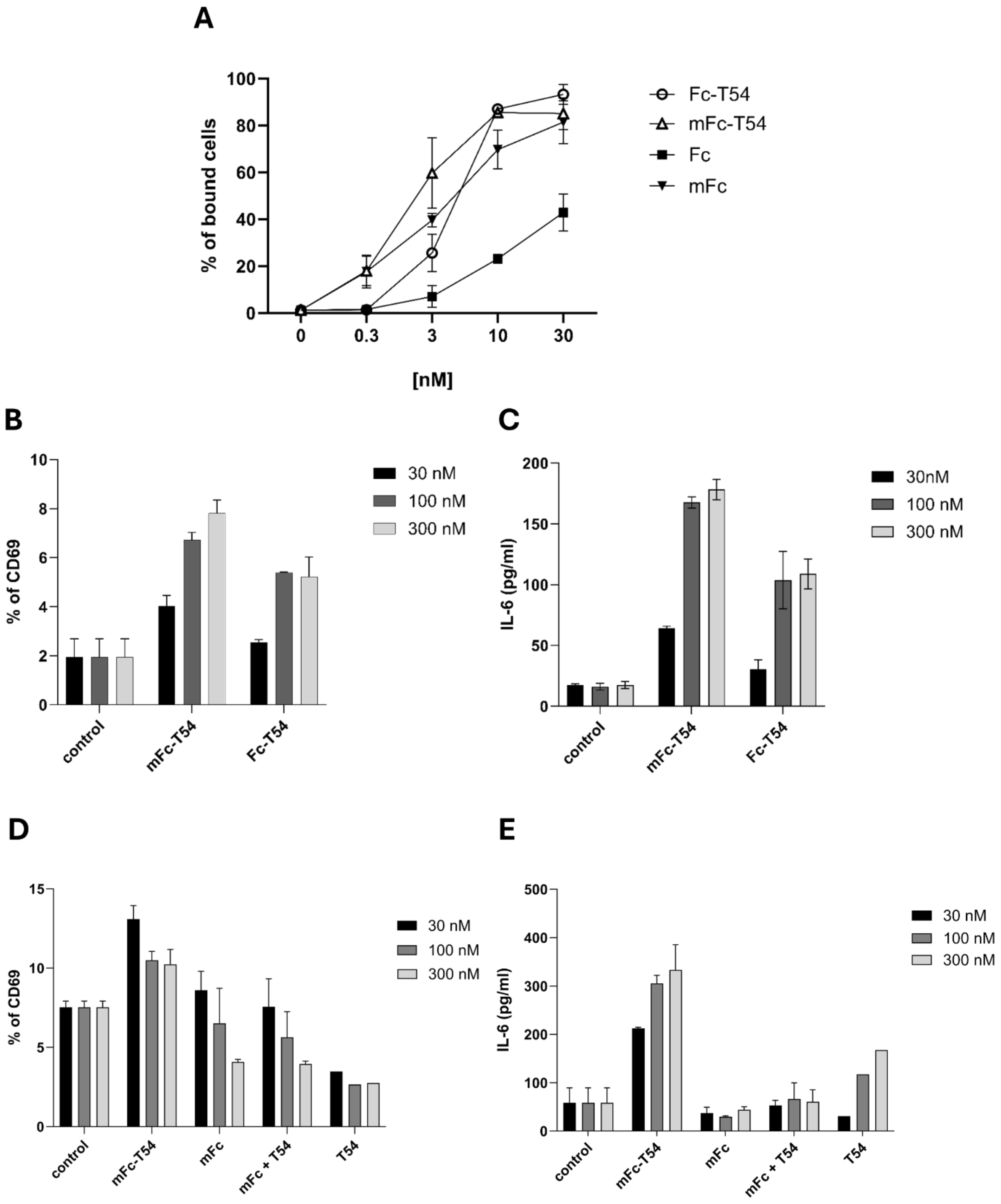


**Figure S2. mFc-T54 ability to bind and activate mouse DCs. A:** mFc-T54, Fc-T54, mFc and Fc binding to DCs. JAWSII DCs were incubated with biotinylated mFc-T54, Fc-T54, mFc and Fc for 30min in 4°C. Then, cells were labeled with fluorescent Streptavidin, and binding was assessed by flow cytometry. The percentage of fluorescent cells was calculated relative to the total number of live cells. **B-E:** mFc-T54- and Fc-T54-mediated activation of DCs, JAWSII cells were incubated with or without mFc-T54, Fc-T54, mFc and Fc. After 24 hours, supernatants were collected to assess IL-6 secretion (**C** and **E**) and cells were labeled with either a fluorescent anti-CD69 Ab (**B** and **D**). The percentage of CD69-positive cells was plotted relative to the number of live cells.
